# Cyclophilin A-mediated *cis/trans* isomerization modulates RIN4 to control intracellular rhizobial infection in legumes

**DOI:** 10.1101/2025.08.11.669618

**Authors:** Takashi Goto, Kasper Røjkjær Andersen, Masaru Bamba, Shusei Sato, Masayuki Sugawara, Kiwamu Minamisawa, Masayoshi Kawaguchi, Jens Stougaard, Yasuyuki Kawaharada

## Abstract

In most legume-rhizobium symbioses, rhizobial colonization occurs through host-derived intracellular infection threads, which enable recruitment of compatible rhizobia while presumably modulating the host immune system to prevent rejection. To investigate how legumes regulate immune responses through post-translational mechanisms during the infection, we focused on Cyclophilin A (CyPA), a peptidyl-prolyl *cis/trans* isomerase. The model legume *Lotus japonicus* encodes three canonical *CyPA* genes. Among them, *LjCyPA1* was characterized through CRISPR/Cas9-mediated knockout analysis and shown to be important for normal intracellular infection of compatible rhizobia. A gain-of-function *LjCyPA1* variant in a soybean cultivar was able to promote symbiosis with not only compatible but also incompatible rhizobia. Structural modeling followed by genetic analysis demonstrated a functional interaction between LjCyPA1 and the immune hub protein LjRIN4. The *cis* conformation of LjRIN4 promoted intracellular rhizobial infection, while the *trans* conformation suppressed it. LjCyPA1 acted with the rhizobial type III secretion system (T3SS) which exhibited a cooperative role between host and symbiont in facilitating infection. Phylogenomic analysis showed that conservation of the CyPA1 orthologue is correlated with the trait of intracellular infection in legumes. Our results contribute to the understanding of how legumes accept symbiotic partners while balancing immune responses.

## Introduction

A fundamental question in symbioses research centers on how plant hosts engage with symbionts. Legume plants interact with nitrogen-fixing bacteria called rhizobia, selectively hosting those that meet specific criteria as beneficial endosymbionts. The legume-rhizobia symbiosis, widely known as root nodule symbiosis, serves as a model system for understanding the host’s decision-making in accepting compatible symbionts. Legumes perceive rhizobial lipochitooligosaccharides (Nod factors) via the cell-surface receptors NFR1/LYK3 and NFR5/NFP (Madsen *et al*., 2003; Radutoiu *et al*., 2003; Limpens *et al*., 2003; Smit *et al*., 2007; Indrasumunar *et al*., 2010, 2011; Broghammer *et al*., 2012). The authentication of rhizobia is further ensured by recognition of bacterial exopolysaccharide (EPS) by the EPR3 receptor (Kawaharada *et al*., 2015, 2017b; Kelly *et al*., 2023). The structure and modifications of Nod factor and EPS are bacteria specific and enable legume-bacteria pairing (López-Lara *et al*., 1996; Roche *et al*., 1996; Skorupska *et al*., 2006; Radutoiu *et al*., 2007; Kawaharada *et al*., 2015). Upon recognizing compatible signals, legumes initiate symbiotic signaling pathway leading to rhizobial infection and nodule formation. Two different modes of infection routes are found in legumes: intracellular and intercellular infection. The intracellular route observed in approximately 75% of legume hosts transports rhizobia from epidermal root-hair cells into cortical cells, via tunnel-like tubular structures called infection threads. In the intercellular route, rhizobia invade through cracks between epidermal cells formed during lateral root emergence (Quilbé *et al*., 2022). Intracellular infection is more efficient than intercellular infection, since the host actively invites rhizobia, contingent upon those rhizobial molecules meeting stricter criteria (Madsen *et al*., 2010; Kawaharada *et al*., 2017b; Acosta-Jurado *et al*., 2019; Montiel *et al*., 2021).

Successful symbiosis requires not only these infection modes but also proper regulation of the immune response that would otherwise reject symbionts. Nod factor perception induces phosphorylation of downstream factors and suppresses the production of reactive oxygen species (ROS) (Shaw & Long, 2003; Liang *et al*., 2013; Rey *et al*., 2019; Feng *et al*., 2021; Wang *et al*., 2025). Cytoplasmic kinases LICK1/2 simultaneously activate symbiotic signaling via trans-phosphorylation of LYK3 and suppress the immune response (Wang *et al*., 2025). These insights highlight the importance of immune suppression for successful symbiosis, yet much is still unknown on how the immune system is modulated during rhizobial infection. Plant immunity consists of two immune systems: microbe/pathogen-associated molecular patterns (MAMPs/PAMPs)-triggered immunity (MTI/PTI) and effector-triggered immunity (ETI). MTI/PTI is a basal immune response triggered by common microbial patterns. To suppress this defense, bacteria deliver effector proteins into host cells, but host plants carrying the corresponding resistance (R) genes recognize these effectors and counterattack with ETI (Jones & Dangl, 2006). Interestingly, rhizobia also attempt to promote infection through effectors known as nodulation outer proteins (Nops) (Marie *et al*., 2001; Cao *et al*., 2017; Miwa & Okazaki, 2017; Piromyou *et al*., 2021; Ma *et al*., 2024), suggesting that these effectors promote symbiosis in the context of compatible interactions. By contrast, when certain rhizobia produce incompatible Nops, the host legume recognizes them and ETI blocks nodulation (Yang *et al*., 2010; Tang *et al*., 2016; Sugawara *et al*., 2018; Zhang *et al*., 2021). Unlike pathogenic interactions, where broad immune responses are typically effective against invaders, symbiosis requires the host legumes to properly modulate immunity depending on whether rhizobia are compatible or incompatible. This balance and immune flexibility support the robustness of the symbiosis system, which serves as a foundation for the host’s environmental adaptability. However, the regulatory factors that enable this immune modulation in the intracellular infection system of legumes remain largely unexplored.

Cyclophilin (CyP) is a peptidyl-prolyl *cis/trans* isomerase that catalyzes the isomerization of proline residues between *cis* and *trans* conformation in target proteins. Among them, Cyclophilin A (CyPA) was first discovered in association with immunity in animal cells (Handschumacher *et al*., 1984). In *Arabidopsis thaliana*, one of the CyPAs (AtCYP18-3, also known as AtROC1), interacts with RIN4 (Li *et al*., 2014), the immune signaling hub involved in both MTI/PTI and ETI (Mackey *et al*., 2002, 2003; Axtell & Staskawicz, 2003; Toruño *et al*., 2018; Ray *et al*., 2019). Intriguingly, AtCYP18-3/AtROC1, particularly its gain-of-function variant, reduces immune response to *Pseudomonas syringae* by suppressing ETI in leaves (Ma *et al*., 2013; Li *et al*., 2014), a function that may be counterproductive for the host. Here, we hypothesize that native CyPAs fulfill a beneficial function in symbiotic interactions. We analyzed homologous CyPAs in the context of root nodule symbiosis using the model legume *Lotus japonicus* and found that LjCyPA1 is required for normal intracellular infection of compatible rhizobia. The gain-of-function *LjCyPA1* variant exhibited enhanced symbiosis with both compatible and incompatible rhizobia. Structural modeling followed by genetic analysis suggested that *cis/trans* isomerization of LjRIN4 influences rhizobial acceptance during intracellular infection, which requires rhizobial effectors. LjCyPA1 is co-conserved with intracellular infection in legumes, and we discuss how the *cis/trans* isomerization of LjRIN4 contributes to acceptance of symbiotic partners during intracellular infection.

## Materials and Methods

### Plant materials and growth conditions

The *L. japonicus* Miyakojima MG-20 ecotype was used as wild-type. The soybean cultivar used in this study was Hardee. Three-day-old MG-20 seedlings were transferred to culture vessels containing sterilized vermiculite with B&D medium and grown for 3 days to adapt. For bacterial growth, *M. loti* MAFF303099 (for *L. japonicus*) was cultured in YEM medium, while *B. diazoefficiens* USDA122 (for soybean) was cultured in HEPES-MES (HM) salt medium (Cole & Elkan, 1973) (supplemented with 0.1% arabinose and 0.025% [wt/vol] of yeast extract). For inoculation, rhizobia were suspended in B&D medium: *M. loti* MAFF303099 in Stock A-D (x1), pH 5.7; and USDA122 in Stock B (x0.5) and Stock C-D (x1) at pH 6.8. DsRed-labeled *M. loti* MAFF303099 were used for microscopic observation.

### CRISPR/Cas9 mutagenesis

To generate *cypA1*, *cypA2*, and *cypA3* knockout plants using a CRISPR/Cas9 system, double-strand DNA oligos were designed using the CRISPR-P program (http://crispr.hzau.edu.cn/CRISPR2/) (Lei *et al*., 2014). These sequences are listed in Table S1. These oligos were cloned into the entry vector pUC_AtU6-oligo (Ito *et al*., 2015). For gRNA expression, an I-SceI-digested gRNA module was inserted into the binary vector pZK _gYSA _FFCas9 (Ito *et al*., 2015). For whole-plant transformation, *Agrobacterium tumefaciens* AGL1 strains were used. Hypocotyl segments from MG-20 wild-type were incubated on papers soaked in co-culture medium (1/10 Gamborg’s B5 salt mixture, 1/10 Gamborg’s vitamin solution, 0.5 µg mL^−1^ BAP, 0.05 µg mL^−1^ NAA, 5 mM MES (pH 5.2), 20 μg ml^−1^ acetosyringone, pH 5.5) containing an AGL1 suspension for 5 days at 24°C in the dark. Next, the segments were transferred onto a callus induction medium (1× Gamborg’s B5 salt mixture, 1× Gamborg’s vitamin solution, 2% sucrose, 0.2 μg ml^−1^ BAP, 0.05 μg ml^−1^ NAA, 10 mM (NH_4_)_2_SO_4_, 0.3% phytagel,12.5 μg ml^−1^ meropen, 15 μg ml^−1^ Hygromycin B, pH 5.5). They were cultured under a 16-hour light/8-hour dark cycle at 24°C and transferred every 5 days for 2–5 weeks. Once the calli turned deep green, they were transferred onto a shoot elongation medium (Gamborg’s B5 salt mixture, Gamborg’s vitamin solution, 2% sucrose, 0.2 μg ml^−1^ BAP, 12.5 μg mL^−1^ meropenem, 0.6% agar, pH 5.5) and grown for 3–6 weeks under the same conditions. The calli were transferred onto a fresh callus induction medium every 7 days. The individual shoots from calli were excised and inserted into a root induction medium (1/2 Gamborg’s B5 salt mixture, 1/2 Gamborg’s vitamin solution, 1% sucrose, 0.5 μg ml^−1^ NAA, 0.9% agar, pH 5.5) and cultivated for 1-2 weeks under the same conditions. Once root formation was evident, they were transplanted into root elongation medium (1/2 Gamborg’s B5 salt mixture, 1/2 Gamborg’s vitamin solution, 1% sucrose, 0.9% agar, pH 5.5) for 2 weeks under the same conditions.

### Construction of ΔT3SS rhizobia

The type III secretion system (T3SS) mutant was constructed using a transconjugation system. Two genomic regions (∼1 kb each) of *M. loti* MAFF303099 and the gentamicin cassette region from pMS246 (Becker *et al*., 1995) were amplified by PCR, and cloned to pK18mob digested by *BamH*I using In-Fusion (Takara Bio Inc.) reaction. This plasmid was transformed into *M. loti* MAFF303099 harboring DsRed via triparental mating with *E. coli* strain MT616 carrying the mobilizing plasmid pRK600 (Finan *et al*., 1986). Subsequently, a double-crossover mutant was selected based on gentamicin resistance and kanamycin sensitivity.

### Hairy root transformation

Hairy root transformation was carried out using *Agrobacterium rhizogenes* AR1193 (for *L. japonicus*) and K599 (for soybean). MG-20 and *cypA1* seedlings, grown for 3 days in darkness followed by 1 day under a 16 h light/8 h dark cycle at 24°C, were excised below the hypocotyls while submerged in an AR1193 suspension containing the appropriate vectors. The seedlings were then co-cultivated on half-strength B5 medium (Wako or Duchefa biochemie) supplemented with 0.02 g L⁻¹ sucrose, 0.5 g L⁻¹ MES, and 0.9% agar at 24°C in darkness for 3 days. Subsequently, they were transferred to a hairy root induction medium (B5 medium with 1% sucrose, Gamborg B5 vitamin solution, 0.5 g L⁻¹ MES, 12.5 μg mL⁻¹ meropenem [Sumitomo Pharmaceuticals or Sigma-Aldrich], and 0.9% agar) and incubated for 10 days under a 16 h light/8 h dark cycle at 24°C. Transgenic hairy roots were confirmed by GFP signal. Plants were inoculated with rhizobia 10 days after being transferred to vermiculite. The soybean cultiver Hardee seedlings were grown in culture vessels containing sterilized vermiculite for 5-7 days in darkness and then excised below the hypocotyls. The cut surfaces were coated with K599, and hairy roots were induced in the culture vessels. After two weeks, non-GFP roots were removed, and plants were transplanted into the culture vessels and inoculated with rhizobia 10 days later.

### RNA sequencing

Roots of MG-20 and the *cypA1* mutant, either without (0 DAI) or with 3 days after inoculation of the *M. loti* ΔT3SS mutant, were harvested. Total RNA was isolated using the RNeasy Plant Mini Kit (QIAGEN), and DNA was removed by DNase treatment (QIAGEN). Libraries were sequenced using an MGI DNBSEQ-T7, generating paired-end reads. All reads were quality-checked by FastQC and adapter trimmed with Trimomatic (ver. 0.33, options: LEADING:20 TRAILING:20 SLIDINGWINDOW:4:15 MINLEN:30). Reads were then mapped to the genome (Gifu v1.2 model) using HISAT2 (ver. 2.1.0). Read counts were obtained using HTSeq (ver. 2.0.7). Normalization and extraction of differentially expressed genes (DEGs) were performed on iDEGES/edgeR-edgeR pipline in the TCC R package.

### Gene expression analysis

Primers used for qRT-PCR are listed in Table S1. Total RNA was isolated from roots using NucleoSpin RNA Plant (MACHEREY-NAGEL). First-strand cDNA was prepared using Maxima H Minus First Strand cDNA Synthesis Kit (Thermo Fisher Scientific). qRT-PCR was performed with LyghtCycler 480 SYBER Green I Master (Roche) on CFX96 Real Time System (BIO-RAD) following the manufacturer’s instructions. Expression of *LjUBQ* was used as a reference.

### Phylogenetic and phylogenomic analysis

Protein and genome sequences obtained from the NCBI Genome database and Phytozome v12 (https://phytozome.jgi.doe.gov/pz/portal.html) were used. Alignment of sequences was performed using Clustal X. To analyze synthetic relationships between *L. japonicus* and multiple plant species, we used the JCVI toolkit (https://github.com/tanghaibao/jcvi). Orthologous gene pairs were identified using the jcvi.compara.catalog ortholog command with protein-coding sequences across 17 species*; Medicago truncatula*, *Trifolium pratense*, *Cicer arietinum*, *Glycine max*, *Glycine soja*, *Vigna angularis*, *Vigna radiata*, *Abrus precatorius*, *Arachis ipaensis*, *Arachis duranensis, Lupinus albus, Lupinus angustifolius*, *Aeschynomene evenia, Fragaria vesca*, *Malus domestica*, *Populus trichocarpa*, and *Arabidopsis thaliana*. Syntenic blocks were detected using MCScan (Wang *et al*., 2012) implemented in JCVI, by comparing *L. japonicus* gene coordinates against synteny anchors, iterating 20 times to refine predictions. For visualization, we followed the JCVI manual guidelines. In the graphical representation, due to polyploidy, only one chromosome was used for *Glycine max*, *Glycine soja*, and *Malus domestica* (e.g., *CyPA1*-associated synteny was observed on chromosomes 4 and 6 of *G. max* and *G. soja*, and on chromosomes 5 and 10 of *Malus domestica*).

### Site-directed mutagenesis of CyPA1 and RIN4

The cording sequences of *LjCyPA1* and *LjRIN4* were amplified from *L. japonicus* MG-20 cDNA. Each PCR fragments were cloned into pENTR/D-TOPO (Invitrogen). *LjCyPA1* variants (gain-of-function: S58F; binding defective: F67A; catalytic defective: H133A) and *LjRIN4* variants (*cis*-conformation mimic: ΔP187, *trans*-conformation mimic: P187V) were generated using the PrimeSTAR Mutagenesis Basal Kit (TAKARA). The primers are listed in Table S1. Along with the original *LjCyPA* and *LjRIN4*, their respective variants in pENTR/D-TOPO, were recombined into the modified Gateway binary vector, proLjUBQ:GW-GFP.

### Protein modeling

AlphaFold2 was used to generate the models of *L. japonicus* CyPA1 and *A. thaliana* CYP18-3/ROC1. No template was used in the modelling process. Structural analysis and figures were made in PyMOL version 3.1.1 (Schrödinger, LLC).

### Microscopy

Fluorescence microscopy was performed with a BX50 upright microscope (Olympus) or with an A1R confocal microscope (Nikon). Images were acquired and analyzed using DP Controller (Olympus) and NIS Elements (Nikon).

## Results

### Identification of CyPA homologues in *Lotus japonicus*

A BLASTP analysis using canonical CyPAs (human CyPA and one of the Arabidopsis CyPAs, AtCYP18-3/ROC1) as queries identified three CyPA proteins in *L. japonicus*, which we named CyPA1 (Lj1g3v3343880), CyPA2 (Lj3g3v3527420), and CyPA3 (Lj3g3v3527430). These CyPAs shared highly conserved amino-acid sequences with AtCYP18-3/ROC1: 92.4% (CyPA1), 92.4% (CyPA2), 84.9% (CyPA3) (Fig. S1a).

AlphaFold2 prediction showed high structural similarity between AtCYP18-3/ROC1 and LjCyPAs with RMSD values of 0.083 Å (CyPA1), 0.106 Å (CyPA2), and 0.101 Å (CyPA3), and conservation of the residues in the active sites (Fig. S1b-d). *CyPA1* is located on chromosome 1, whereas *CyPA2* and *CyPA3* are tandemly positioned on chromosome 3 (Fig. 1a). According to our RNA-seq (Goto *et al*., 2022), these mRNAs were abundant in both *Mesorhizobium loti*-inoculated and non-inoculated roots in early-infection stage (0-3 days after inoculation [DAI]) (Fig. S2).

**Figure 1.**
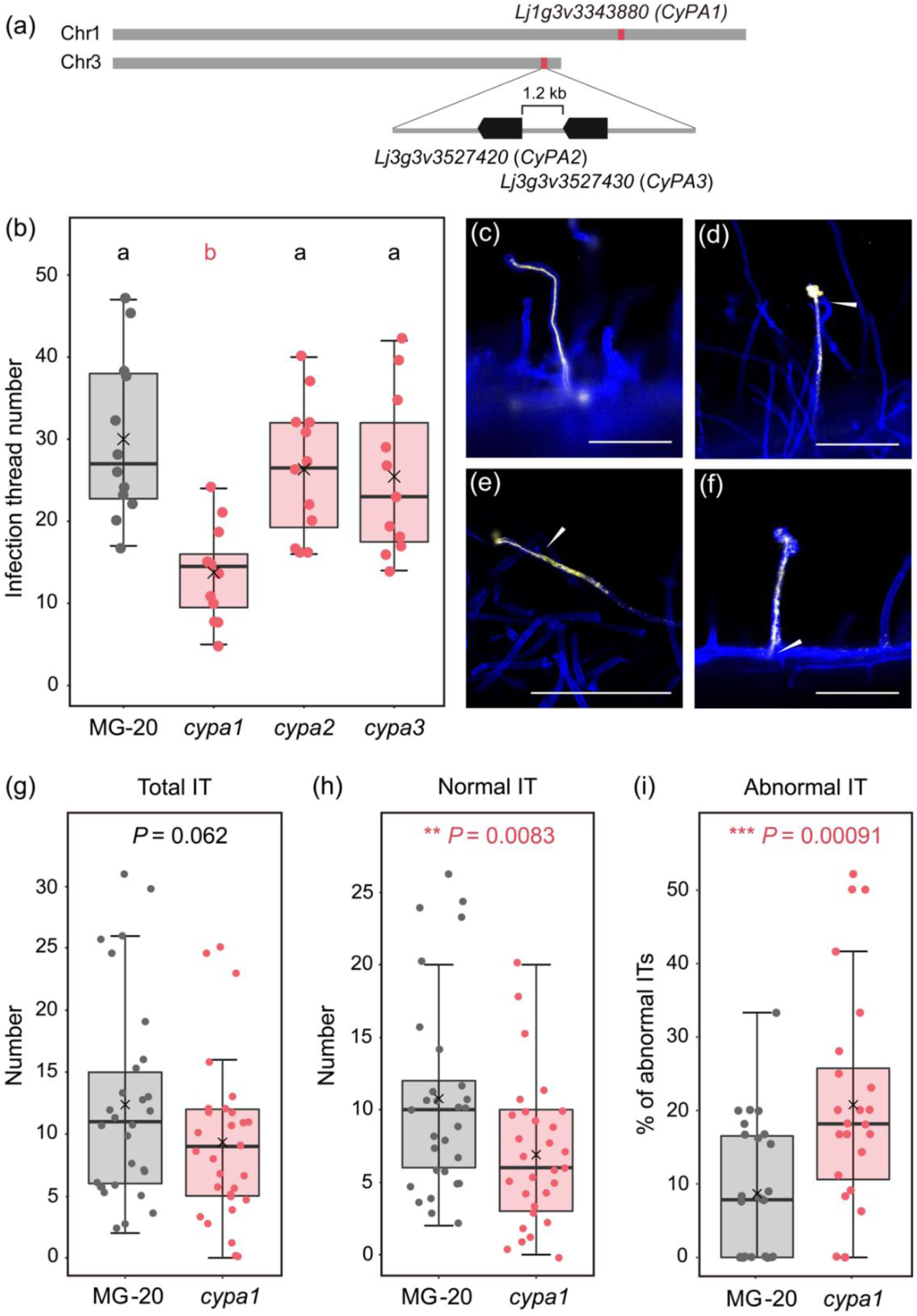
*CypA1* ensures the process of rhizobial intracellular infection. **a** Genomic location of *CyPA1* (*Lj1g3v3343880*) on chromosome 1, and *CyPA2* (*Lj3g3v3527420*), and *CyPA3* (*Lj3g3v3527430*) on chromosome 3. **b** Number of infection threads (ITs) in MG-20 wild-type, *cypA1*, *cypA2*, and *cypA3* mutants at 8 days after inoculation. ANOVA followed by Tukey’s honestly significant difference (HSD) test (*P* < 0.05). Different letters indicate significant differences. **c-f** Representative phenotypes of normal ITs in MG20 wild-type (c) and abnormal ITs in the *cypA1* mutant (d-f) in each stage of IT elongation: initiation (d), middle elongation (e), or elongated ITs (f), and *M. loti* was released within root hairs from the tip of ITs. Arrowheads indicate the tip of each IT. Scale bar: 100 µm. **g-i** The number of total ITs (normal + abnormal) (g) and normal ITs (h), and the proportion of abnormal ITs (number of abnormal ITs / total ITs) (i) at 5 DAI. Asterisks indicate statistically significant differences (g-h: Welch’s t-test and i: Fisher’s exact test).

### CyPA1 facilitates intracellular root-hair entry of symbiotic bacteria

We utilized CRISPR technology to create knockout lines with nonsense mutations in *CyPA1*, *CyPA2*, and *CyPA3* genes of *L. japonicus* MG-20 wild-type (Fig. S3). Among these, *CyPA1* mutation had a significant impact on nodule symbiosis with the highly compatible rhizobium *M. loti* MAFF303099. The *LjcypA1* mutant plants showed a decrease in the number of normal infection threads (ITs) at 8 DAI compared to MG-20 wild-type and the other *LjcypA2* and *LjcypA3* mutant plants (Fig. 1b). The infection phenotype of *LjcypA1* was validated using an independent *LjcypA1* mutant (*LjcypA1-2*) (Fig. S4) and was rescued by constitutive expression of *LjCyPA1* using a hairy root transformation system (Fig. S5). Compared to the normal ITs in MG-20 (Fig. 1c), ITs in the *LjcypA1* mutant were frequently abnormal: IT progression was arrested at initiation or elongation, and rhizobia were released within root hairs (Fig. 1d-f). When we examined the infection phenotypes at an earlier timepoint (5 DAI), the total number of ITs did not differ (Fig. 1g), but abnormal ITs were significantly more frequent in the *LjcypA1* mutant (Fig. 1h-i). These findings indicate that *LjCyPA1* serves ensuring effective IT progression within root hairs.

### CyPA1 and rhizobial T3SS cooperate in rhizobial infection and nodulation

Rhizobia use type III secretion system (T3SS) to deliver effector proteins (T3Es) into the host cells to regulate symbiosis (Marie *et al*., 2001; Miwa & Okazaki, 2017). To explore the relationship between LjCyPA1 and rhizobial T3Es during root-hair infection, we engineered a mutant of *M. loti* MAFF303099 lacking the T3SS genes (ΔT3SS) (Fig. 2a). At the early infection stage (5 DAI), the *LjcypA1* mutant showed significantly reduced normal ITs in total ITs after inoculation with the *M. loti* ΔT3SS mutant (Fig. 2bc). 16% of ITs in MG-20 were abnormal following the *M. loti* ΔT3SS mutant inoculation, a slight increase from the 8% observed with *M. loti* MAFF303099 (Fig. 2d). In contrast, the *LjcypA1* mutant exhibited 53% abnormal ITs following the *M. loti* ΔT3SS mutant inoculation, a dramatic increase from the 21% observed with *M. loti* MAFF303099 (Fig. 2d).

**Figure 2.**
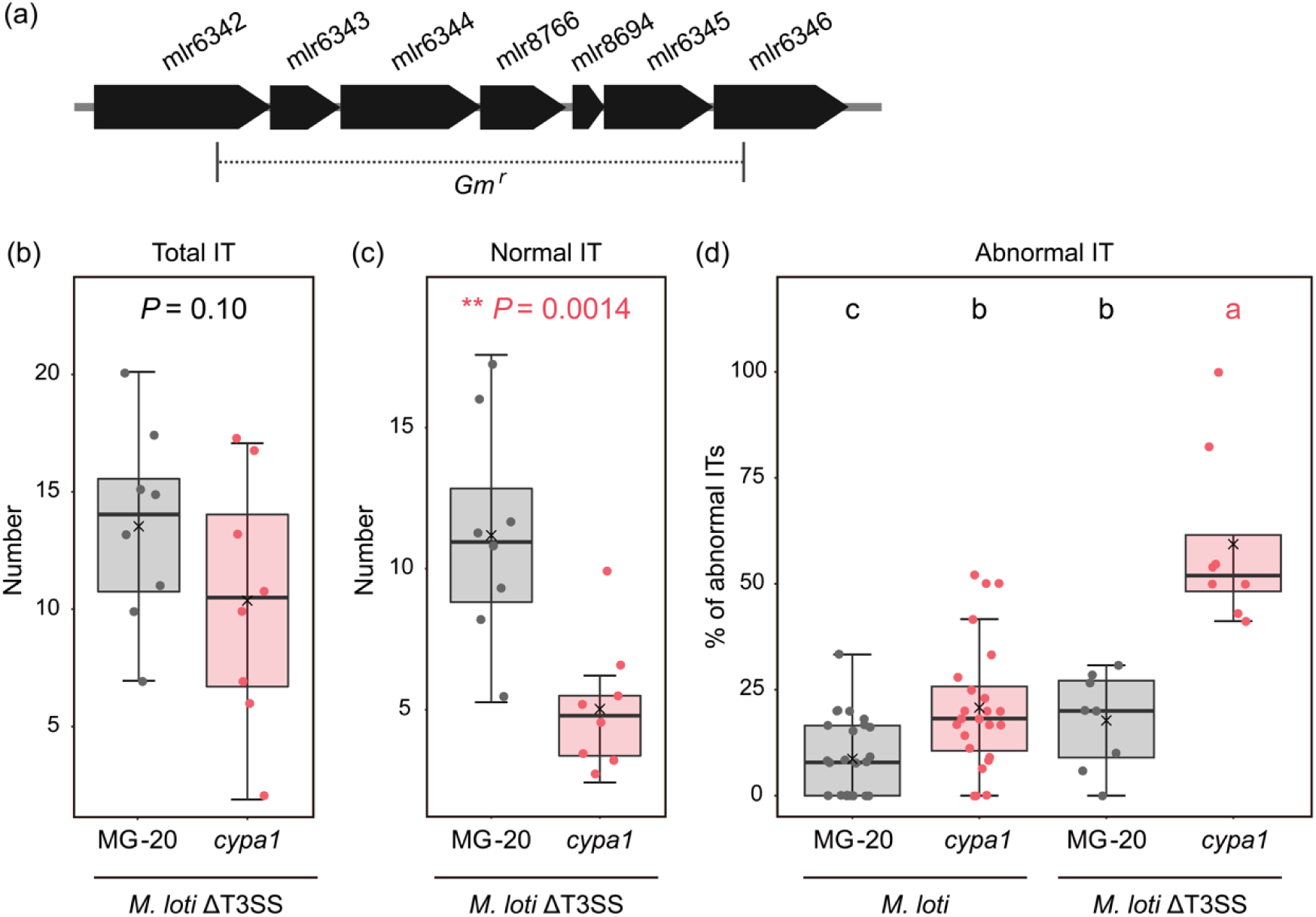
CyPA1 and compatible rhizobial T3SS facilitate intracellular infection. **a** T3SS-deficient *M. loti* MAFF303099 (the *M. loti* ΔT3SS mutant). **b-d** The number of total ITs (normal + abnormal) (b) and normal ITs (c), and the percentage of abnormal ITs (number of abnormal ITs / total ITs) (d) at 5 DAI with *M. loti* MAFF303099 or the *M. loti* ΔT3SS mutant. Asterisks and different letters indicate that differences are statistically significant (b, c, e: Welch’s t-test and d: Fisher’s exact test).

In nodule organogenesis, there were morphological differences that could affect their maturity (Fig. 3). Brownish-white nodule primordia with black wounds were only observed in the *LjcypA1* mutant two weeks after inoculation (Fig. 3b and d). This result showed that the *LjcypA1* mutant increased nodulation with abnormal morphology not only in epidermal infection. On the other hand, transgenic hairy roots showed individual loss of *CyPA1* or inoculation with the *M. loti* ΔT3SS mutant did not affect nodule number; however, the simultaneous loss of both *CyPA1* and T3SS severely impaired nodule formation (Fig. 3e and f). These results indicate that host CyPA1 and rhizobial T3Es redundantly function in the compatible symbiosis.

**Figure 3.**
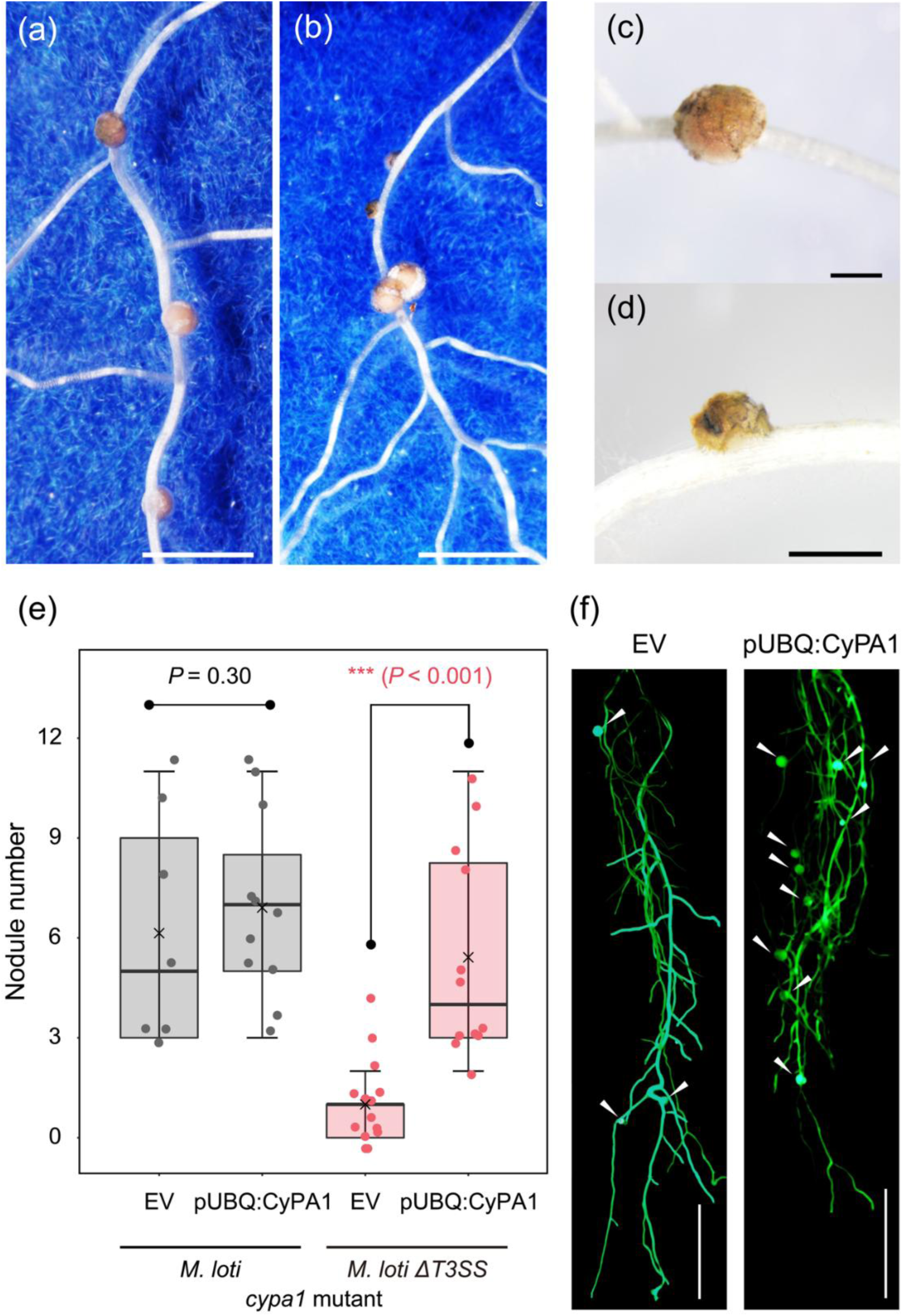
Nodulation phenotypes in MG-20 and the *cypA1* mutant with the *M. loti* ΔT3SS mutant. **a** and **c** Nodules on MG-20 wild-type at 14 days after inoculation (DAI). **b** and **d** Brownish-white nodule primordia on the *cypA1* mutant at 14 DAI. Scale bars: 500 µm (a-b), 100 µm (c-d). **e** The nodule number in hairy roots of the *cypA1* mutants harboring EV (empty vector with GFP) and *pUBQ:CyPA1-GFP* vector 3 weeks after inoculation. Asterisks indicate statistical difference by Welch’s t-test. **f** Representative phenotype of hairy roots of *cypA1* mutants harboring EV (Left) and *pUBQ:CyPA1-GFP* vector (Right) with the *M. loti* ΔT3SS mutant. Nodules are indicated by arrowheads. (Scale bars, 1 cm.)

### CyPA1 and rhizobial T3SS affect immune and symbiotic expression

To investigate transcriptomic differences associated with host genotype (MG-20 vs. the *LjcypA1* mutant) and plant responses to the *M. loti* ΔT3SS mutant (non-inoculation vs. 3 DAI), we conducted RNA sequencing. A total of 356 differentially expressed genes (DEGs) were identified (FDR < 0.05; Table S2), including several resistance and defense-related genes. Quantitative RT-PCR was then used to further compare expression patterns following non-inoculation, *M. loti* MAFF303099 inoculation, and the *M. loti* ΔT3SS-inoculation in MG-20 and the *LjcypA1* mutant (Fig. S6). In MG-20 roots, the *Lotus* homologue of *RPS2*, an NBS-LRR class resistance gene known to suppress pathogen infection in *Arabidopsis* leaves (Mindrinos *et al*., 1994; Bent *et al*., 1994), was down-regulated following inoculation with *M. loti* MAFF303099, but not with the *M. loti* ΔT3SS mutant. While in the *LjcypA1* mutant, no such down-regulation was observed with either mesorhizobial strain (Fig. S6). These results suggest that the expression of *RPS2* is modulated by both LjCyPA1 and rhizobial T3Es. In contrast, a *Chitinase* gene was down-regulated after inoculation with both mesorhizobial strains in MG-20, while its expression remained stable in the *LjcypA1* mutant (Fig. S6), indicating CyPA1-dependent but not T3E. Interestingly, when the *Ljcypa1* mutant was inoculated with the *M. loti* ΔT3SS mutant, a *Defensin-like* gene resulted in dramatic induction (Fig. S6), suggesting that CyPA1 and T3E components synergize in its expression. We also examined the expression pattern of several symbiotic genes. *ERN1*, a symbiotic transcription factor for intracellular rhizobial infection (Kawaharada *et al*., 2017a; Yano *et al*., 2017; Cerri *et al*., 2017; Montiel *et al*., 2021), was not induced in the *cypA1* mutant upon inoculation with the *M. loti* ΔT3SS mutant (Fig. S6). Meanwhile, *RPG*, a regulator of polarized IT elongation (Arrighi *et al*., 2008; Li *et al*., 2023; Lace *et al*., 2023), showed no significant induction after inoculation with the *M. loti* ΔT3SS mutant compared to *M. loti* MAFF303099 inoculation in MG-20 (Fig. S6). *NFR1*, whose expression is reduced following compatible rhizobia inoculation (Frank *et al*., 2023), was also down-regulated in the *Ljcypa1* mutant when inoculated with both *M. loti* MAFF303099 and the *M. loti* ΔT3SS mutant (Fig. S6). Together, these results demonstrate that CyPA1 influences the expression of both immune-related and symbiotic genes involved in IT formation during the early stages of infection, through both T3E-dependent and -independent mechanisms.

### Gain-of-function CyPA1 enhances symbiosis with both compatible and incompatible rhizobia

A gain-of-function mutant of CyPA (*CyPA^S58F^*; *roc1* mutation) was previously identified in Arabidopsis (Ma *et al*., 2013), and has been shown to strongly suppress immunity against pathogens (Li *et al*., 2014). To investigate whether a gain-of-function LjCyPA1 affects common signaling components also involved in the symbiotic nodulation, we examined the effect on intracellular infection and nodule formation. Constitutive expression of *LjCyPA1^S58F^*significantly increase the number of ITs and nodules after inoculation with *M. loti* MAFF303099 compared to controls expressing either an empty vector or native *LjCyPA1* (Fig. 4a and b). When inoculated with the *M. loti* ΔT3SS mutant, *LjCyPA1^S58F^* also promoted nodule formation, although to a lesser extent compared to when inoculated with *M. loti* MAFF303099 (Fig. S7).

**Figure 4.**
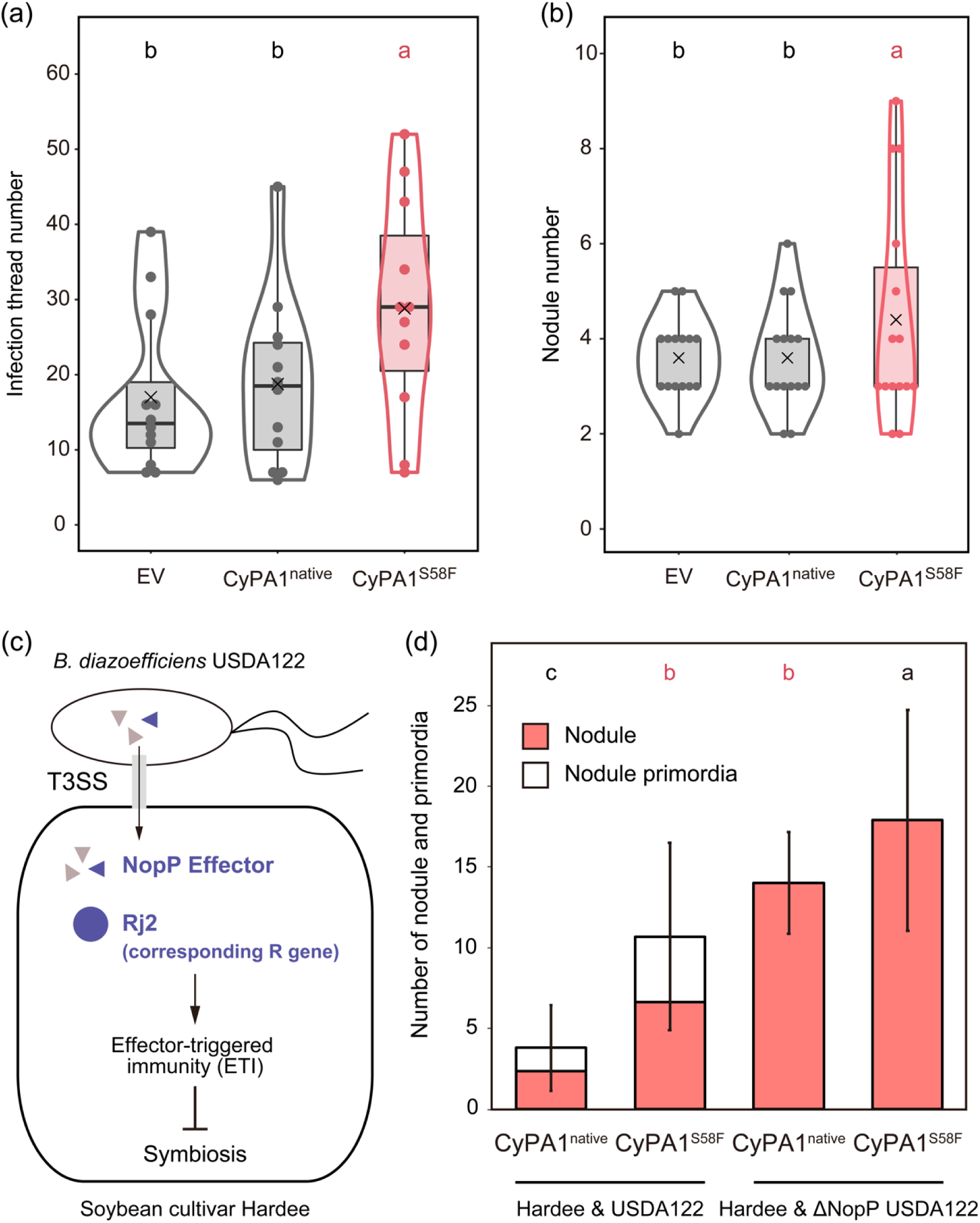
Gain-of-function of *CyPA1* promotes symbiosis with both compatible and incompatible rhizobia. a-b. The number of infection threads (a) and nodules (b) in hairy roots of MG-20 harboring EV (control-1), *pUBQ:LjCyPA1-GFP* vector (control-2), and *pUBQ:LjCyPA1^S58F^-GFP* vector (gain-of-function). Infection threads and nodules were counted 2 and 3 weeks after inoculation, respectively. **c** A schematic illustration of effector-triggered immunity (ETI) in interaction with incompatible *Bradyrhizobium diazoefficiens* USDA122 with soybean cultivar Hardee. Hardee’s Rj2 recognizes NopP effector secreted by *B. diazoefficiens* USDA122 and activates ETI, and suppresses nodule formation. **d** The number of nodules (pink) and primordia (white) in hairy roots of soybean cultivar Hardee with expressing *pUBQ:LjCyPA1-GFP* and *pUBQ:LjCyPA1^S58F^-GFP*. Error bars indicate means ± SDs (n > 10 hairy roots). ANOVA followed by Tukey’s HSD test (*P* < 0.05). Different letters indicate significant differences.

Building on these findings, we explored whether the gain-of-function mutation alleviates severe symbiotic incompatibility caused by effector-triggered immunity (ETI). We focused on the interaction between the soybean cultivar Hardee and *Bradyrhizobium diazoefficiens* USDA122. The host R gene *Rj2* recognizes the incompatible NopP effector and triggers immunity (Fig. 4c) (Sugawara *et al*., 2018, 2019). In this incompatible *B. diazoefficiense* USDA122 inoculation, native *LjCyPA1* expression in Hardee could not promote nodulation, resulting in very few or no nodules (Fig. 4d). However, the gain-of-function *LjCyPA1^S58F^*significantly promoted nodulation (Fig. 4d). Furthermore, NopP-deficient mutant (USDA122ΔNopP), which reverts to a compatible strain, showed high nodule formation with the gain-of-function *LjCyPA1^S58F^* expression roots (Fig. 4d), similar to *M. loti* MAFF303099 inoculation in *LjCyPA1 ^S58F^* variant in MG-20 (Fig. 4b). These results indicate that a gain-of-function of *LjCyPA1* promotes symbiotic nodulation regardless of host-rhizobia compatibility.

### A *cis/trans* isomerization of RIN4 is required for root hair infection

AtCYP18-3/ROC1 interacts with and catalyzes *cis/trans* isomerization of RIN4, a plant immune signaling hub (Li *et al*., 2014). *L. japonicus* possess a single LjRIN4 (Lj3g3v0730080) and our structural modeling predicted that LjCyPA1 forms a complex with an LjRIN4 peptide (182–KGAAV<u>P</u>KFGEWD–193), in which proline-187 interacts with the hydrophobic pocket of LjCyPA1 (Fig. 5a). In this model, a π–π stacking interaction between phenylalanine-67 of LjCyPA1 and phenylalanine-189 of LjRIN4 was predicted to be important for this binding (Fig. 5a). Additionally, histidine-133 of LjCyPA1 was expected to interact with proline-187 of LjRIN4 to facilitate its isomerization (Fig. 5a). To investigate the importance of these aromatic ring interactions in nodule symbiosis, we replaced phenylalanine-67 with Alanine (F67A) or Histidine-133 with Alanine (H133A) in LjCyPA1. As a result, neither LjCyPA1^F67A^ (binding defective mutant) nor LjCyPA1^H133A^ (catalytic defective mutant) could complement nodulation in the *LjcypA1* mutant (Fig. 5b). These results indicate that LjCyPA1-mediated isomerization and binding to LjRIN4 are required for nodule symbiosis. Furthermore, we investigated the impact of *cis*/*trans* isomerization of RIN4 on nodule symbiosis. Proline peptide bonds are naturally biased towards the *trans* form, but *cis/trans* isomerases increase the proportion of the *cis* form. We induced stable *cis* and *trans* isomers (LjRIN4 ^ΔP187^ and LjRIN4 ^P187V^, respectively as described previously (Li *et al*., 2014)) in hairy roots of MG-20 and inoculated with *M. loti* MAFF303099. The LjRIN4^ΔP187^ transgenic roots promoted nodule and normal IT formation (Fig. 5c and d). In contrast, LjRIN4^P187V^ did not affect nodule formation but inhibited normal IT formation (Fig. 5c and d). The opposite phenotypes of RIN4^ΔP187^ and RIN4^P187V^ indicate that shifting the equilibrium from *trans* to *cis* is crucial for symbiosis. Interestingly, when inoculated with the *M. loti* ΔT3SS mutant, RIN4^ΔP187^ did not promote normal IT and nodule formation (Fig. 5c and d). These findings indicate that unknown T3Es in *M. loti* MAFFF303099 are required for promoting symbiosis, working in coordination with the *cis/trans* isomerization at the proline-187 of LjRIN4.

**Figure 5.**
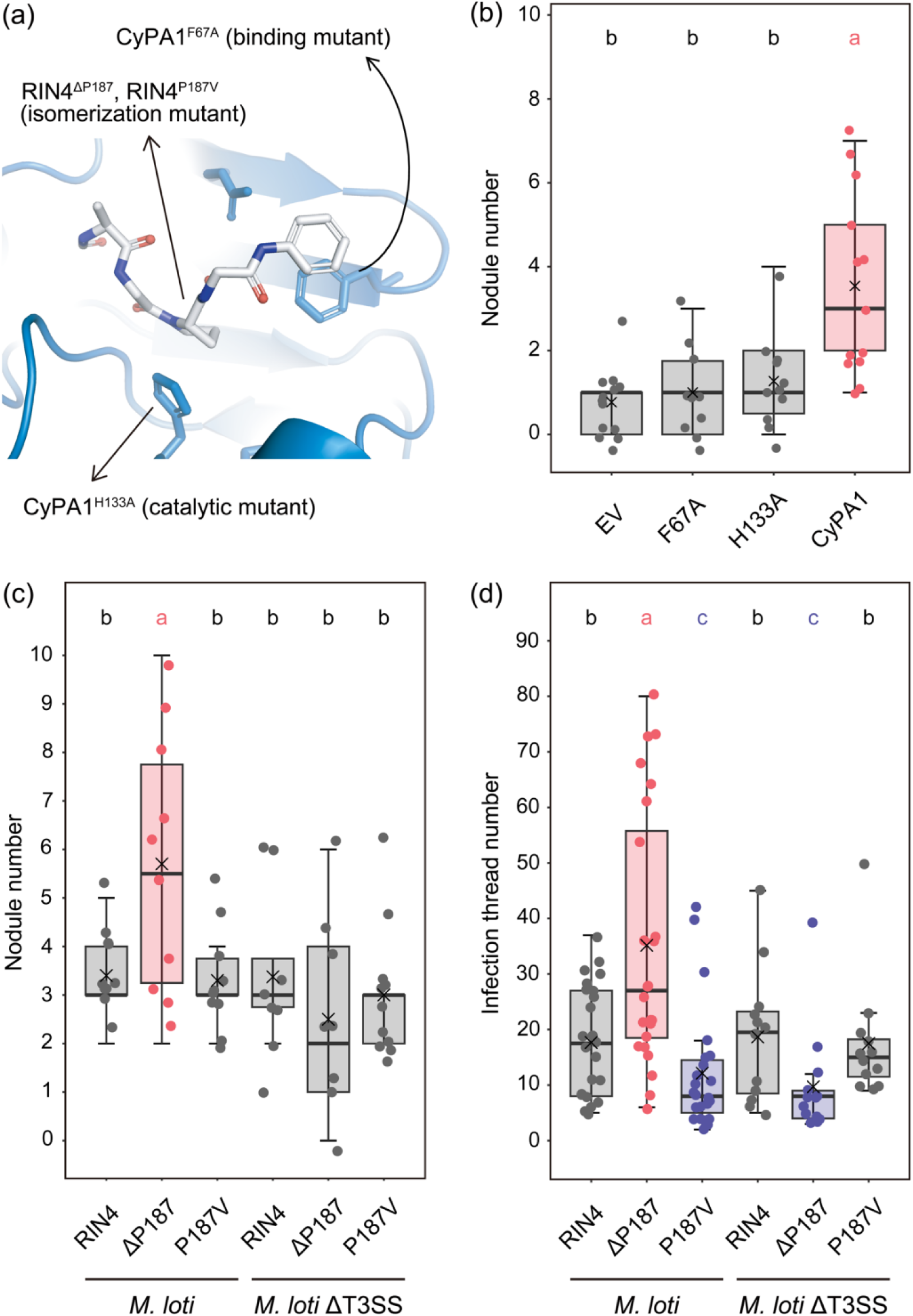
CyPA1-catalyzed *cis/trans* isomerization is essential for symbiosis. **a** Interaction sites between CyPA1 and RIN4 peptide (182-KGAAVPKFGEWD-193) *in silico*. **b** The nodule number in hairy roots of *cypA1* mutants harboring EV (negative control), *pUBQ:LjCyPA1^F67A^-GFP* vector (binding mutant), *pUBQ:LjCyPA1^H133A^-GFP* vector (catalytic mutant), and *pUBQ:LjCyPA1-GFP* vector (positive control), 3 weeks after inoculation with the *M. loti* ΔT3SS mutant. **c-d** The number of nodules (c) and infection threads (d) in hairy roots of MG-20 with constitutive expression of RIN4 (control), RIN4^ΔP187^ (*cis* conformer), RIN4^P187V^ (*trans* conformer). Infection threads and nodules were counted 2 and 3 weeks after inoculation with *M. loti* MAFF303099 or the *M. loti* ΔT3SS mutant, respectively. ANOVA followed by Tukey’s HSD test (*P* < 0.05). Different letters indicate significant differences.

### Loss of CyPA1 correlates with absence of intracellular infection in basal legumes

To investigate the evolutionary conservation of the *CyPA1*-mediated control of intracellular infection, we performed a genome-wide analysis of legumes and non-legumes. Previous phylogenetic analysis has shown that plant CyPs have undergone repeated duplications (Singh *et al*., 2020); however, the evolutionary trajectory of specific CyPAs have not been examined. We performed a synteny analysis of the genomic regions surrounding LjCyPA1 orthologues loci in legumes and non-legumes with high-quality genome. LjCyPA1 orthologues were well conserved in non-legumes as well as legumes (Fig. 6). However, notably, LjCyPA1 orthologues were absent in some early-branching legume genera, such as *Lupinus* (*albus*, *angustifolius*) and *Arachis* (*duranensis*, *ipaensis*) that do not form intracellular ITs (Fig. 6), suggesting the evolutionary loss of CyPA1 in these genera. As an exception, another basal legume *Aeschynomene evenia* possesses a *CyPA1* orthologue, suggesting the CyPA1 was independently lost in the *Lupinus* and *Arachis* genera (Fig. 6). In contrast, LjCyPA2/CyPA3 orthologues were conserved in those species (Fig. S8). These data demonstrate an evolutionary link between CyPA1 and symbiotic traits and suggest that *CyPA1* is evolutionarily important for intracellular infection in legumes that develop ITs.

**Figure 6.**
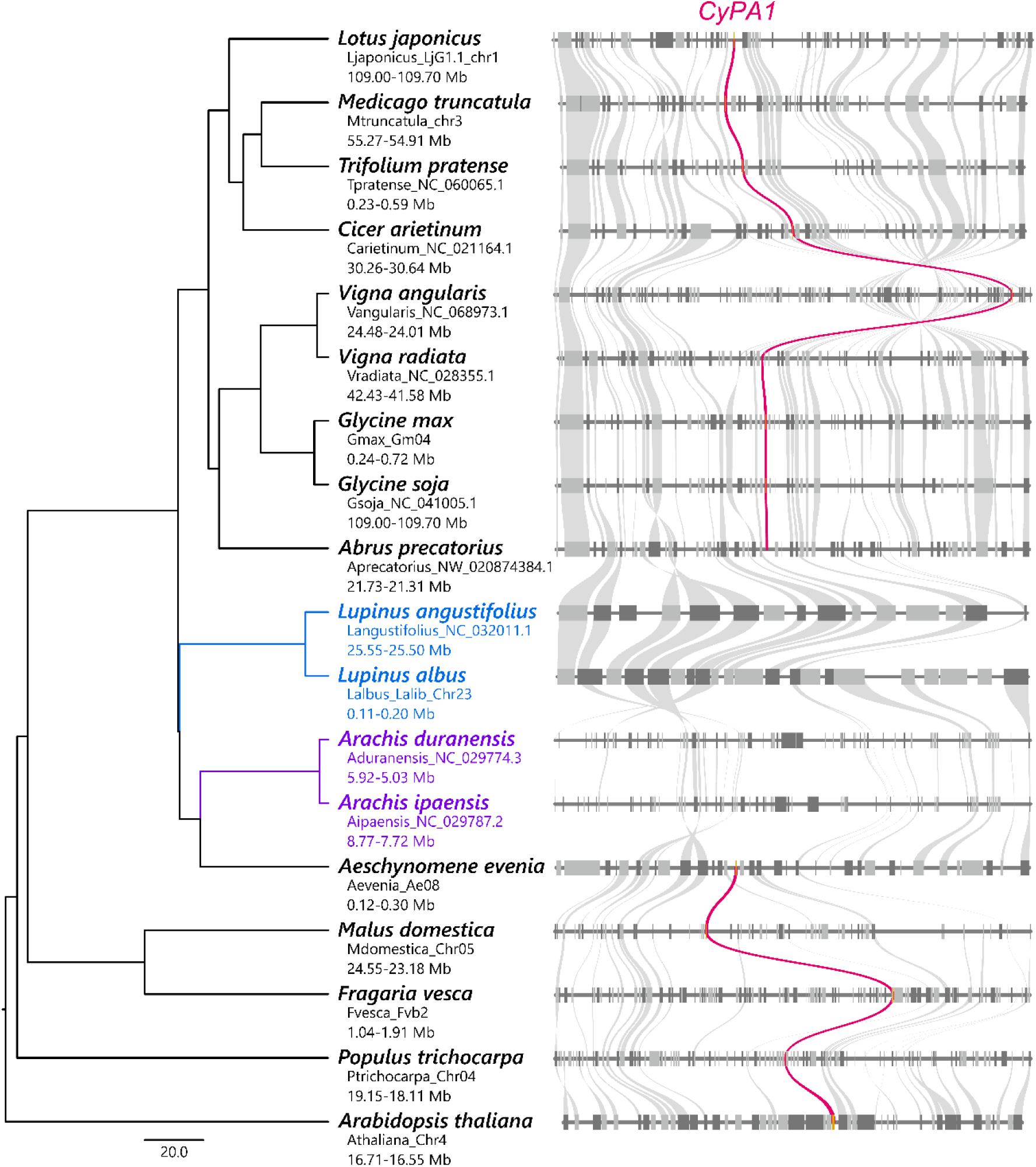
*CyPA1* conservation in representative legumes and non-legumes. Orthologous genes in each specific block are connected by lines of gray or pink colors. *CyPA1* orthologue is highlighted in pink. Dark gray represents genes on the minus strand, while light gray represents genes on the plus strand. The phylogenetic tree to the left of the synteny blocks was obtained from TimeTree5, with the scale bar indicating divergence time in million years ago (Mya).

## Discussion

Among post-translational modifications, peptidyl-prolyl *cis/trans* isomerization is a universal process that alters protein conformation that can switch the direction of signaling. A wide range of studies, including structural biology and reaction kinetics, have underscored the importance of Cyclophilins (CyPs) in catalyzing this isomerization (Li & Cui, 2003; Fanghänel & Fischer, 2004; Lu *et al*., 2007; Jakob & Schmid, 2009; Hamelberg & McCammon, 2009; Camilloni *et al*., 2014). One of the CyPs, Cyclophilin A (CyPA) has been extensively studied in animals since its discovery as a binding protein for the immunosuppressive drug (Handschumacher *et al*., 1984). In contrast, the plant CyPA has undergone duplication and diversification, limiting our understanding of its physiological function (Singh *et al*., 2020). A decade ago, forward genetics in Arabidopsis led to the identification of *roc1*, a gain-of-function mutant of one of the CyPAs, shedding light on the *cis/trans* isomerization in plants. Subsequent studies revealed that AtCYP18-3/ROC1 catalyzes the *cis/trans* isomerization of RIN4, a key signaling hub in plant immunity (Li *et al*., 2014). However, it remained a puzzling aspect of plant-microbe interactions how CyPA play a paradoxical role in suppressing host immunity against pathogens. In this study, we investigated the role of CyPA in the context of root nodule symbiosis. We found that *LjCyPA1*, one of the *L. japonicus CyPAs*, has a native function in facilitating intracellular infection of compatible rhizobia (Fig. 1 and 2). A gain-of-function variant, LjCyPA1^S58F^, accelerated symbiosis even with otherwise incompatible *B. diazoefficiens* USDA122 in a soybean cultivar Hardee (Fig. 4d), highlighting the importance of *cis/trans* isomerization for the proper regulation of host immune responses.

Phenotypic analysis, combined with structural modeling of catalysis and binding, show that LjCyPA1-catalyzed *cis/trans* isomerization is required for symbiosis (Fig. 5a and b). Engineered stable *cis* and *trans* conformers of LjRIN4 at Proline-187 induced opposite phenotypes (Fig. 5c and d), indicating that RIN4’s conformational state plays a role in determining the acceptance of symbionts. Interestingly, this effect depends on the presence of T3SS in *M. loti* MAFF303099, as the promotion of infection through *cis* isomerization at Proline-187 was observed only when unknown T3Es were present (Fig. 5c and d). On the other hand, LjCyPA1 and T3SS exhibit synergistic effects on infection and nodulation, as their simultaneous loss of both causes a stronger phenotypic effect than the loss of either one alone (Fig. 2 and 3). Together, these results suggest that LjCyPA1 promote rhizobial infection through two distinct mechanisms: one involves a T3E-independent pathway, such as down-regulation of *Chitinase* (Fig. S6). The other involves LjCyPA1 acting on *cis* isomerization of LjRIN4, which requires compatible rhizobial T3Es for its function (Fig. 5c and d). This function was supported by *RPS2* and *Defensin-like* expression patterns (Fig. S6). Thus, while CyPA1 and T3Es act synergistically at the overall level, a hierarchical relationship exists in which T3E-dependent regulation occurs at the level of specific targets. Previous studies of the plant immune system have shown that various pathogenic T3Es, such as AvrRpm1, AvrB, AvrRpt2, HopF2, and HopZ3, directly interact with RIN4 (Mackey *et al*., 2002, 2003; Axtell & Staskawicz, 2003; Wilton *et al*., 2010; Lee *et al*., 2015). Among them, AvrRpm1 and AvrB effector phosphorylate AtRIN4 and activate RPM1-mediated defense response (Mackey *et al*., 2002). Also, a novel phosphorylation site of RIN4, unique to the nitrogen-fixing clade, has been recently identified as essential for nodule symbiosis (Tóth *et al*., 2023). Given these findings, both *cis/trans* isomerization and phosphorylation through unknown rhizobial compatible T3Es may alter RIN4 conformation, influencing symbiosis. Understanding how RIN4 adopts distinct conformational states under various microbial influences will be an important direction for future research. RIN4 may serve as a molecular switch integrating symbiotic and immune signals via multiple post-translational modifications.

Given that the evolutionary robustness of intracellular infection system is linked to the CyPA1 orthologue conservation in legumes (Fig. 6), the biological significance of CyPA1’s function in balancing the immune response may be related to the advantage of intracellular infection in legumes. Intracellular infection via ITs represents a “closed” infection system and is superior in that legume actively recruits compatible rhizobia while modulating immunity to avoid rejection. This study demonstrates that LjCyPA1 serves a fundamental role in this pathway by preventing IT abortion during elongation (Fig. 1). Early-branching legumes that do not provide ITs have lost CyPA1 (Fig. 6), which may reflect evolutionary pressures to mitigate risks such as hijacking of root hairs by pathogen. Alternatively, in these lineages, the CyPA1 loss may have gradually weakened the dominance of the IT strategy, eventually leading to the loss of ITs. Our study provides key insights into how CyPA1-mediated *cis/trans* isomerization balances the immune response, ensuring robust symbiotic interactions.

## Acknowledgements

We thank Ms. Sachiko Tanaka (National Institute for Basic Biology; NIBB) and Dr. Kamolchanok Umnajkitikorn (Suranaree University of Technology) for their technical suggestions. We also thank Dr. Mitsutaka Fukudome (Kagoshima University), Dr. Jesús Montiel (Universidad Nacional Autónoma de México), Dr. Simona Radutoiu (Aarhus University), Dr. Stig Uggerhøj Andersen (Aarhus University), Dr. Johan Quilbé (Aarhus University) and Dr. Aleksandr Gavrin (Aarhus University) for constructive discussions. This work was supported by Optics and Imaging Facility (NIBB), Data Integration and Analysis Facility (NIBB), Grants-in-Aid for Scientific Research (23K19376), and the JSPS Overseas Research Fellowship (202460148).

## Supporting Information

**Supplementary figure 1. Sequential and structural alignment of LjCyPAs and AtROC1.** a Amino acid sequences of full-length AtROC1 (At4g38740), LjCyPA1 (Lj1g3v3343880), LjCyPA2 (Lj3g3v3527420), and LjCyPA3 (Lj3g3v3527430). Important residues for the catalytic function are highlighted in asterisks. b-d Structural alignment of each LjCyPA with AtROC1. Superposition of each LjCyPA (blue) and AtROC1 (orange) structural models with a root-mean-square deviation (RMSD). Important residues for the catalytic function are highlighted in b.

**Supplementary figure 2.** mRNA abundance of *LjCyPA1*, *LjCyPA2*, and *LjCyPA3* in early infection stage. MG-20 (wild-type; light-gray) and *daphne* (mutant which shows excessive infection of rhizobia; dark-gray) at 0 (non-inoculation), 1, 2, and 3 DAI. Error bars indicate means ± SDs. (n = 20 roots for each biological replicate).

**Supplementary figure 3.** Schematic illustration of CRISPR/Cas9-induced nonsense mutation in *LjCyPAs*. Each gRNA induces the insertion or deletion of several base pairs, causing a frameshift. The black arrowhead indicates the site where the insertion or deletion occurs, and the pink color represents the newly formed stop codon.

**Supplementary figure 4. The number of infection threads in MG-20 wild-type and *cypA1* another mutant allele *(LjcypA1-2*).** Each dot represents the number of infection threads of each plant. n = 10 (MG-20 and *cypA1-2*). Asterisks indicate that differences are statistically significant (Welch’s t test).

**Supplementary figure 5. Infection thread numbers in *LjcypA1* mutant are restored in hairy roots harboring *LjCyPA1* expression.** Each dot represents the number of infection threads of each plant in control (empty vector; EV) and constitutive expression of *LjCyPA1* (*pUBQ:CyPA1-GFP*). n = 12 (EV and *pUBQ:CyPA1-GFP*). Asterisks indicate that differences are statistically significant (Welch’s t test).

Supplementary figure 6. Quantitative RT-PCR analysis of immune-/defense-related and symbiotic gene expression in MG-20 and the *cypA1* mutant, with or without *M. loti* MAFF303099 or its ΔT3SS mutant. MG-20 (left) and the *cypA1* mutant (right) under mock conditions (non-inoculated, white) or 3 days after inoculation with *M. loti* MAFF303099 (light gray) or its ΔT3SS mutant (dark gray). Error bars indicate the mean ± SD. (n = 10 roots per biological replicate). ANOVA followed by Tukey’s HSD test (P < 0.05). Different letters indicate statistically significant differences. Asterisks indicate notable changes in gene expression.

**Supplementary figure 7. Gain-of-function of *LjCyPA1* promotes symbiosis with *M. loti* MAFF303099 and the *M. loti* ΔT3SS.** The number of nodules in MG-20 hairy roots harboring *pUBQ:CyPA1-GFP* vector (control; gray) and *pUBQ:CyPA1^S58F^-GFP* vector (pink) 3 weeks after inoculation. Asterisks indicate statistical difference by Welch’s t-test.

**Supplementary figure 8. *CyPA2* and *CyPA3* conservation in representative legumes and non-legumes.** Orthologous genes in each specific block are connected by lines of gray or pink colors. *CyPA2/3* orthologue is highlighted in pink. Dark gray represents genes on the minus strand, while light gray represents genes on the plus strand. The phylogenetic tree to the left of the synteny blocks was obtained from TimeTree5, with the scale bar indicating divergence time in million years ago (Mya).

**Supplementary table 1. List of primers**

**Supplementary table 2. Differential gene expression in RNA-seq**

## Contributions

T.G. conceived the project. T.G., Y.K., and K.R.A. designed the experiments. Y.K. created the ΔT3SS mutant of *M. loti* MAFF303099. K.R.A. conducted the structural analysis of CyPA1 and RIN4. T.G. and M.B. performed the synteny analysis. T.G. performed all other experiments. K.M. and M.S. provided the soybean cultivar Hardee, as well as *B. diazoefficiens* USDA122 and its ΔNopP mutant. M.K. and J.S. provided the experimental environments. T.G. and Y.K. wrote the manuscript with feedback from K.R.A., S.S., M.K., and J.S.

