## Supplementary Information for "Cyclophilin A-mediated *cis/trans* isomerization modulates RIN4 to control intracellular rhizobial infection in legumes"

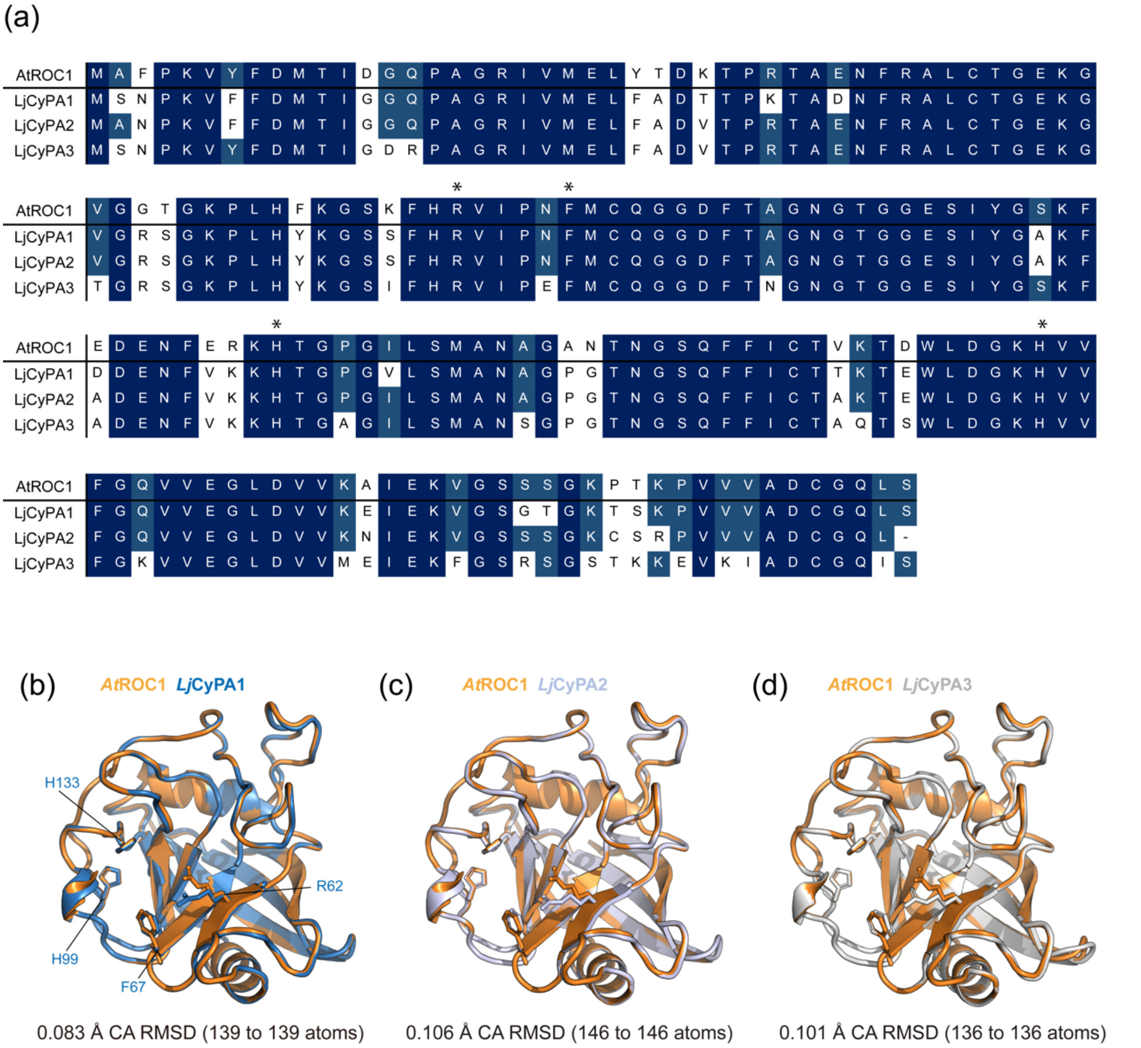


**Supplementary figure 1. Sequential and structural alignment of LjCyPAs and AtROC1. a** Amino acid sequences of full-length AtROC1 (At4g38740), LjCyPA1 (Lj1g3v3343880), LjCyPA2 (Lj3g3v3527420), and LjCyPA3 (Lj3g3v3527430). Important residues for the catalytic function are highlighted in asterisks. **b-d** Structural alignment of each LjCyPA with AtROC1. Superposition of each LjCyPA (blue) and AtROC1 (orange) structural models with a root-mean-square deviation (RMSD). Important residues for the catalytic function are highlighted in b.

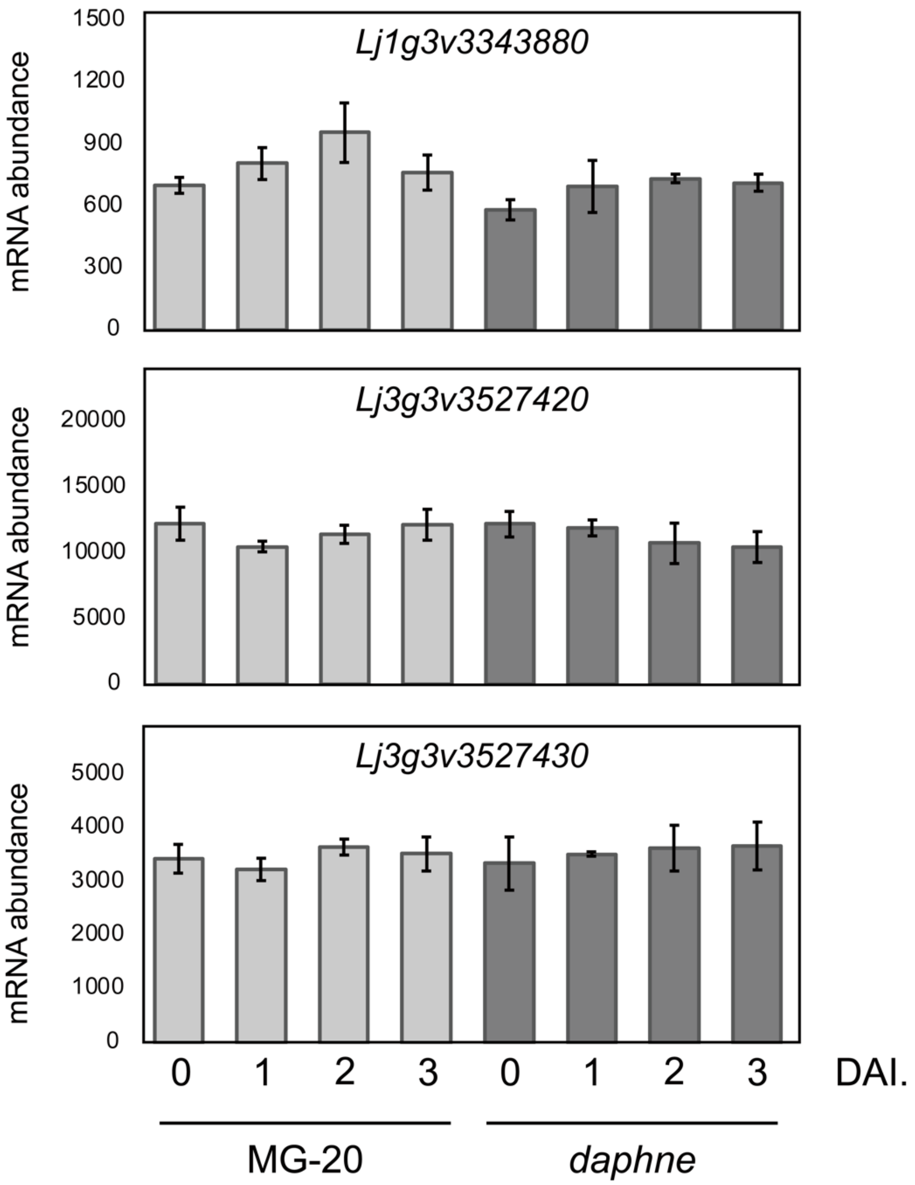


**Supplementary figure 2. mRNA abundance of *LjCyPA1*, *LjCyPA2,* and *LjCyPA3* in early infection stage.** MG-20 (wild-type; light-gray) and *daphne* (mutant which shows excessive infection of rhizobia; dark-gray) at 0 (non-inoculation), 1, 2, and 3 DAI. Error bars indicate means ± SDs. (*n = 20 roots* for each biological replicate).

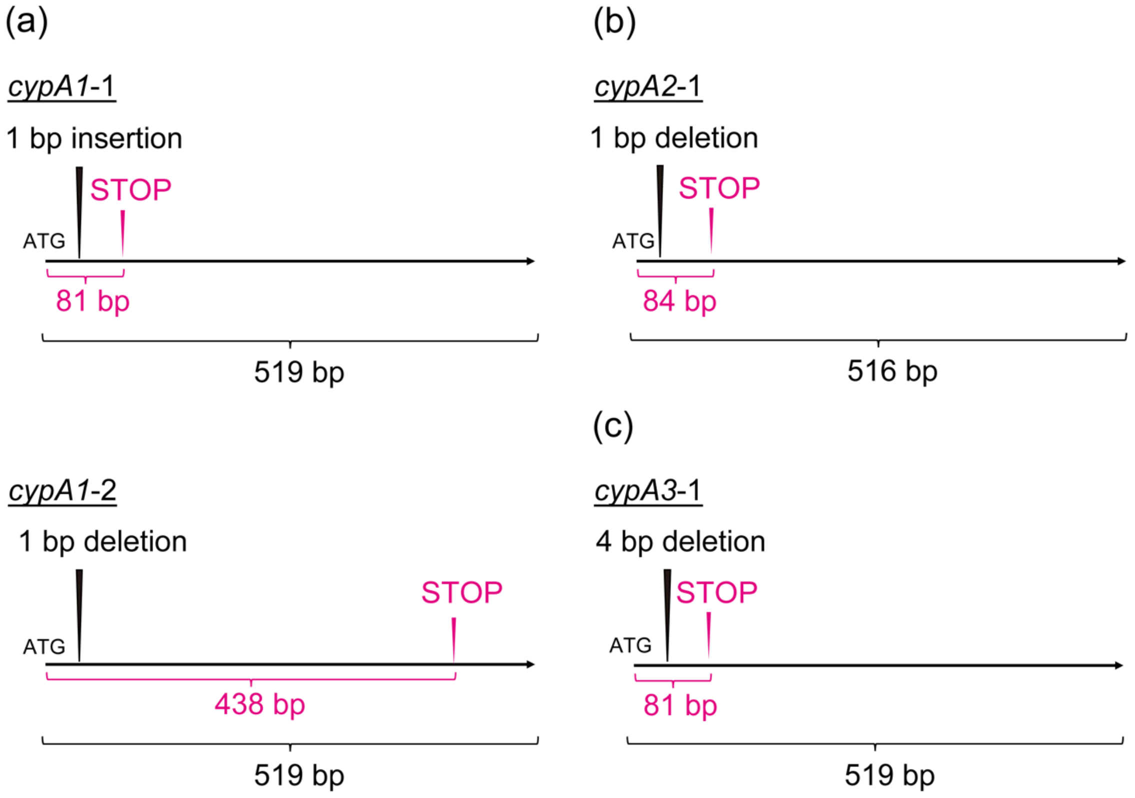


**Supplementary figure 3. Schematic illustration of CRISPR/Cas9-induced nonsense mutation in *LjCyPAs*.** Each gRNA induces the insertion or deletion of several base pairs, causing a frameshift. The black arrowhead indicates the site where the insertion or deletion occurs, and the pink color represents the newly formed stop codon.

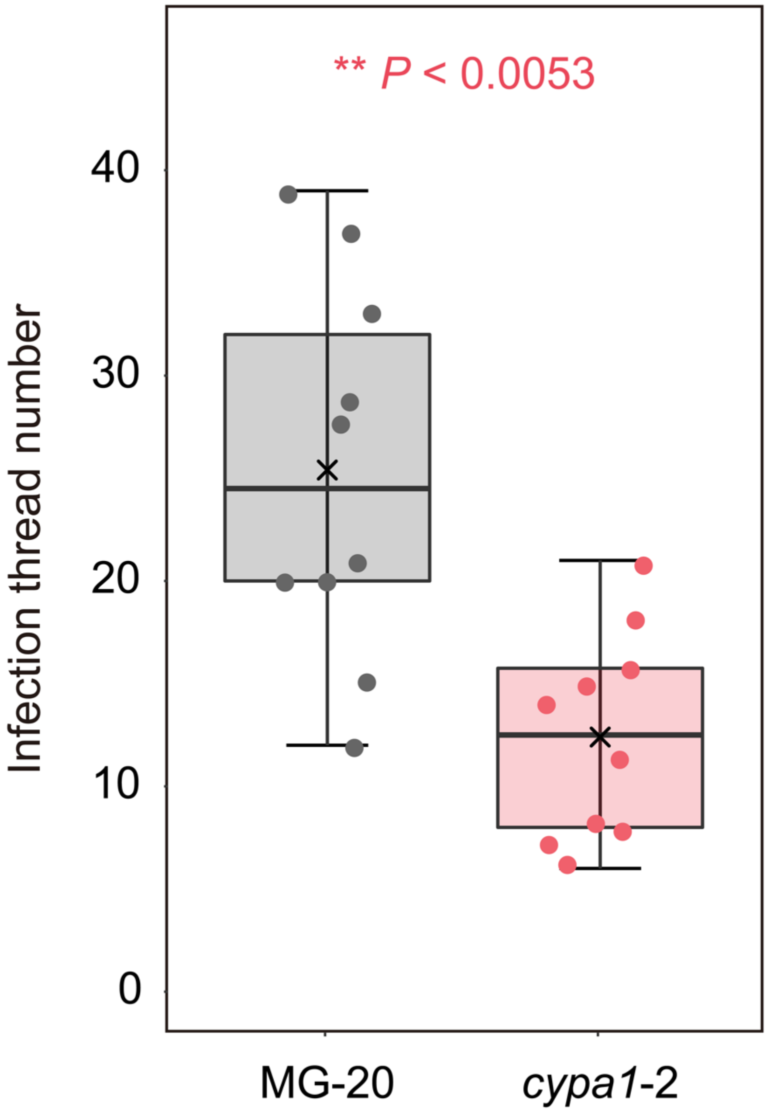


**Supplementary figure 4. The number of infection threads in MG-20 wild-type and *cypA1* another mutant allele (Lj*cypA1-2*).** Each dot represents the number of infection threads of each plant. *n* = *10* (MG-20 and *cypa1*-2). Asterisks indicate that differences are statistically significant (Welch's t test).

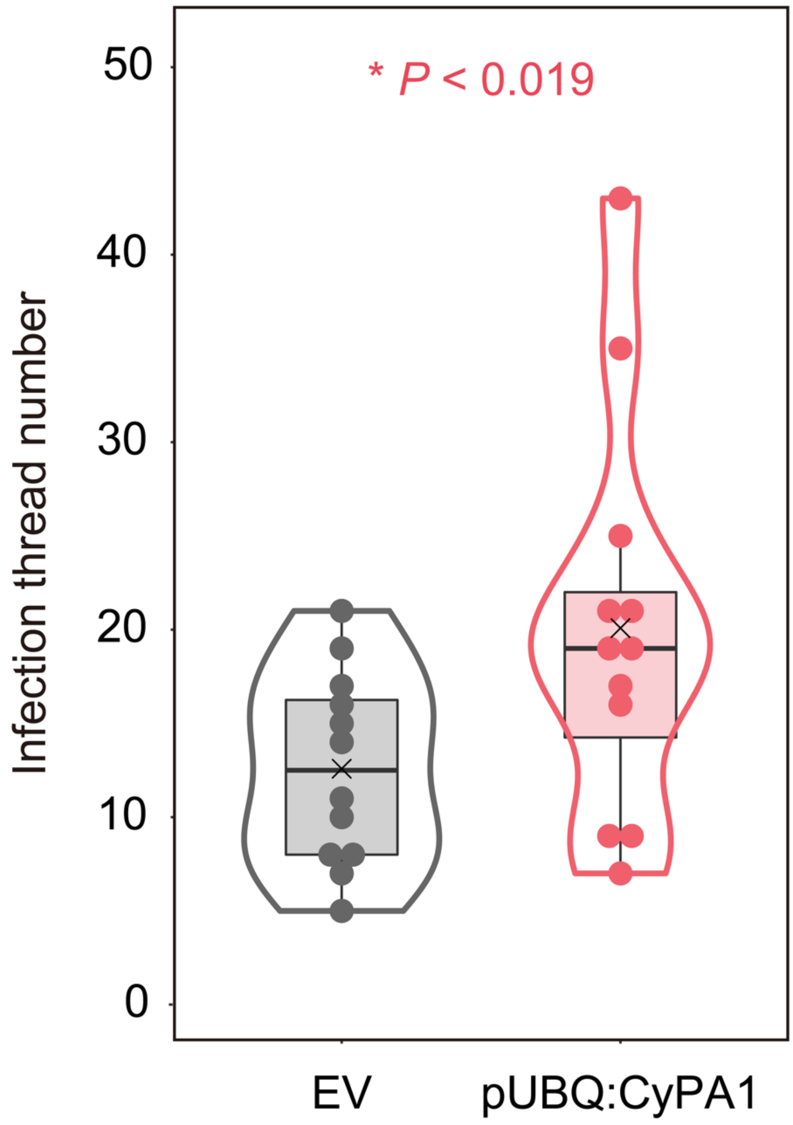


**Supplementary figure 5. Infection thread numbers in *LjcypA1* mutant are restored in hairy roots harboring *LjCyPA1* expression.** Each dot represents the number of infection threads of each plant in control (empty vector; EV) and constitutive expression of *CypA1* (pUBQ:CyPA1). $n = 12$ (EV and pUBQ:CyPA1). Asterisks indicate that differences are statistically significant (Welch's t test).

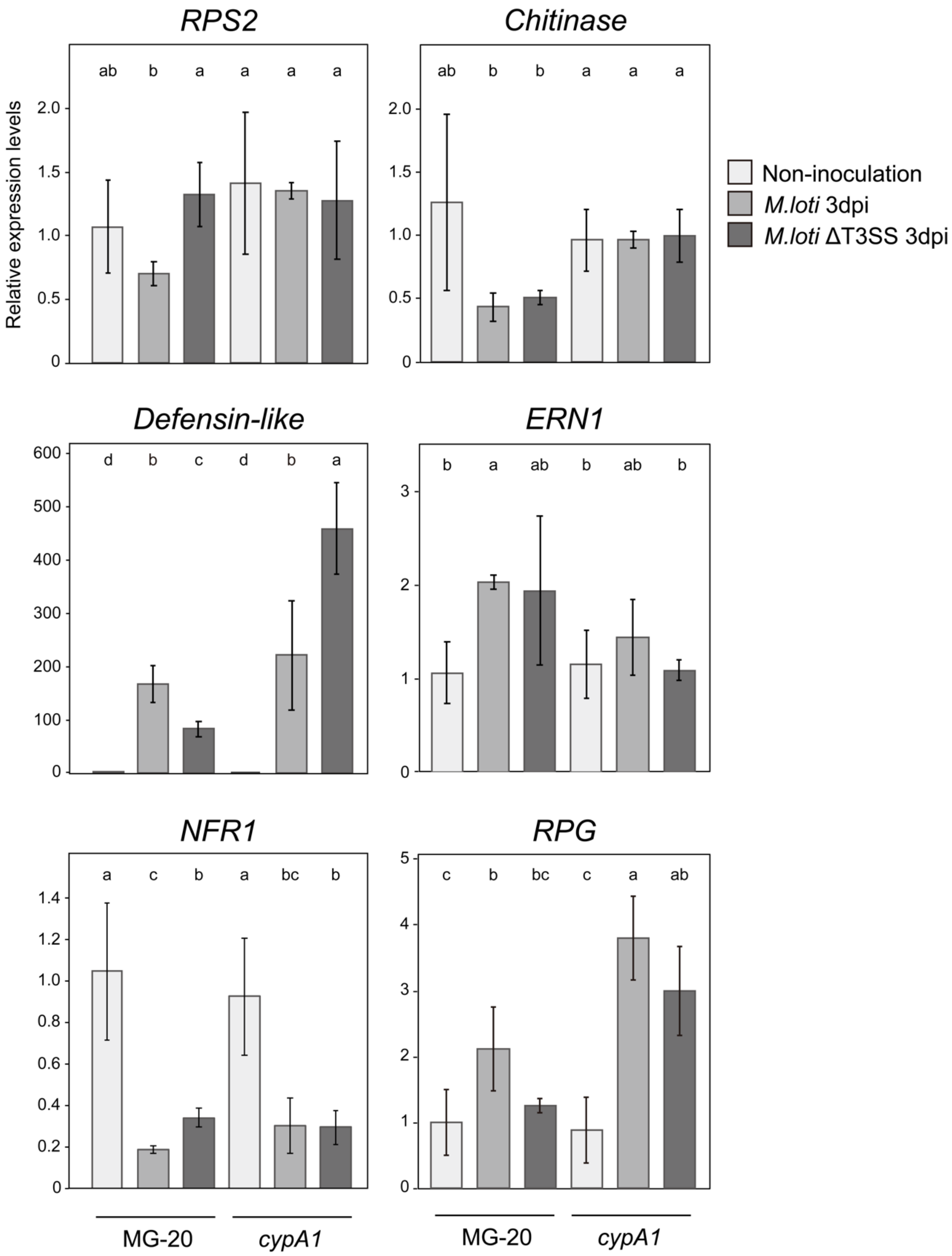


**Supplementary figure 6. Quantitative RT-PCR analysis of immune-/defense-related and symbiotic gene expression in MG-20 and the *cypA1* mutant, with or without *M.***

***loti* MAFF303099 or its ΔT3SS mutant.** MG-20 (left) and the *cypA1* mutant (right) under mock conditions (non-inoculated, white) or 3 days after inoculation with *M. loti* MAFF303099 (light gray) or its ΔT3SS mutant (dark gray). Error bars indicate the mean ± SD. (*n = 10 roots* per biological replicate). ANOVA followed by Tukey's HSD test ($P < 0.05$). Different letters indicate statistically significant differences. Asterisks indicate notable changes in gene expression.

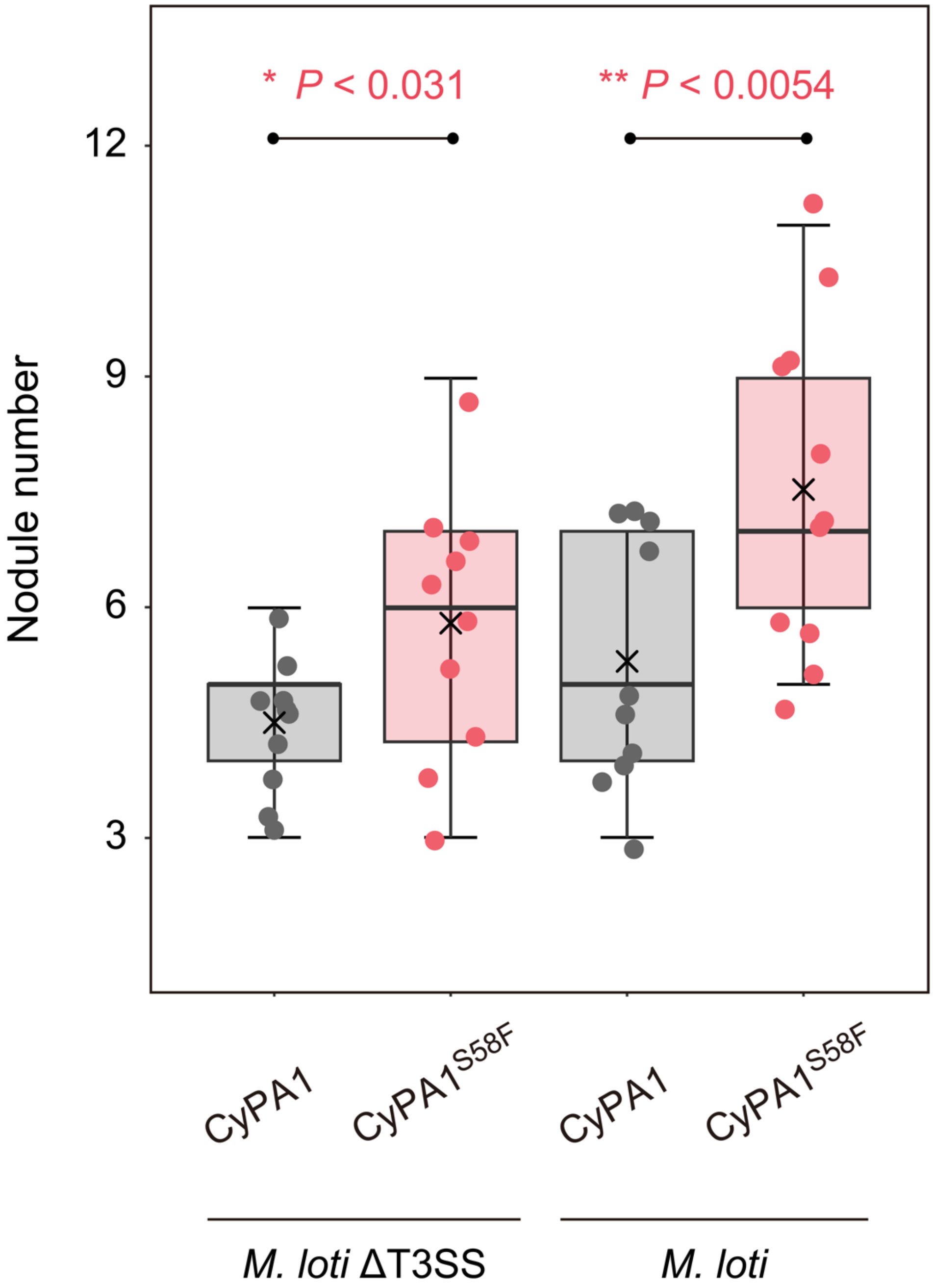


**Supplementary figure 7. Gain-of-function of *LjCyPA1* promotes symbiosis with *M. loti* MAFF303099 and the *M. loti* ΔT3SS.** The number of nodules in MG-20 hairy roots harboring *pUBQ:CyPA1-GFP* vector (control; gray) and *pUBQ:CyPA1*$^{S58F}$*-GFP* vector (pink) 3 weeks after inoculation. Asterisks indicate statistical difference by Welch's t-test.

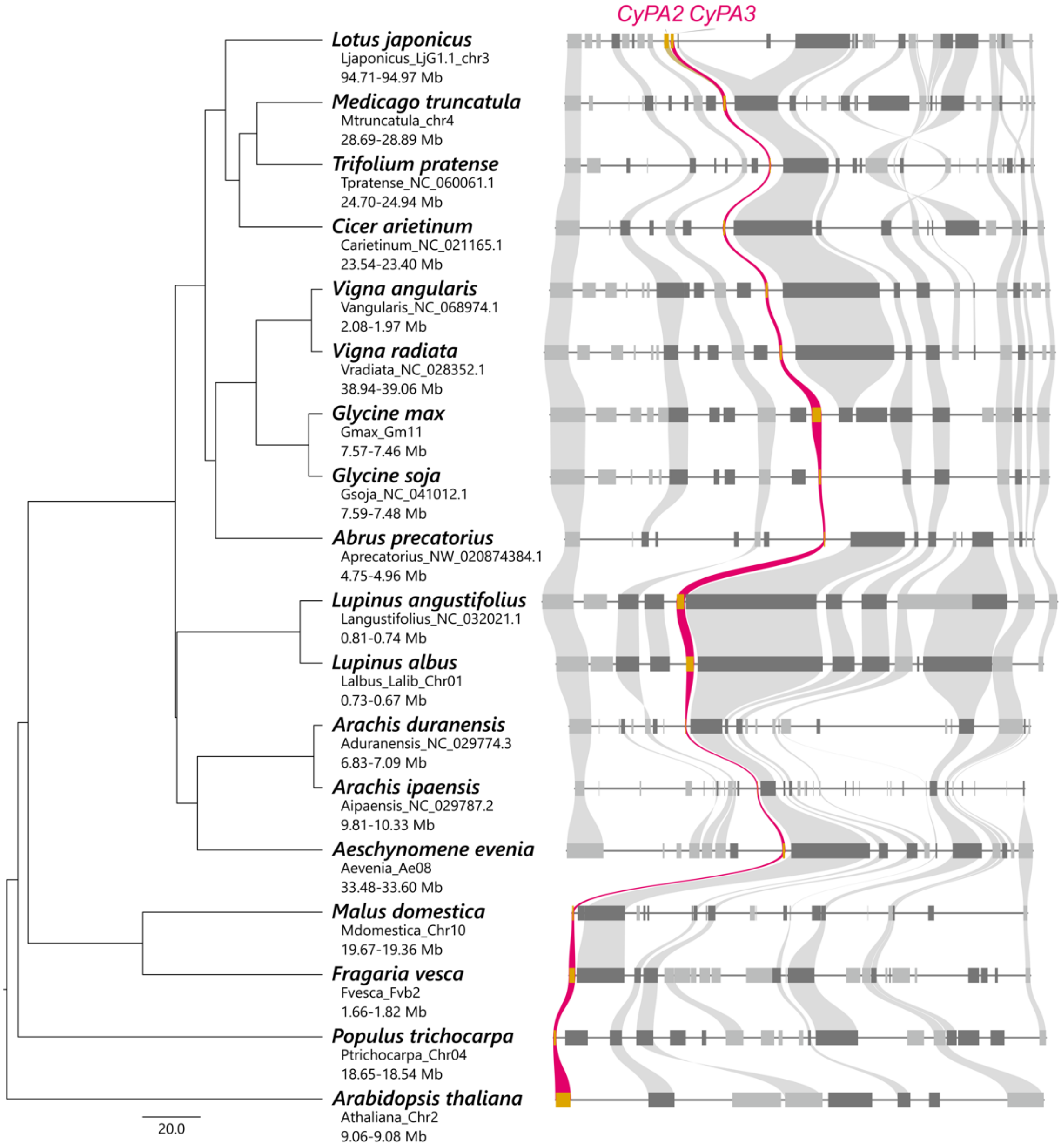


**Supplementary figure 8. *CyPA2* and *CyPA3* conservation in representative legumes and non-legumes.** Orthologous genes in each specific block are connected by lines of gray or pink colors. *CyPA2/3* orthologue is highlighted in pink. Dark gray represents genes on the minus strand, while light gray represents genes on the plus strand. The phylogenetic tree to the left of the synteny blocks was obtained from TimeTree5, with the scale bar indicating divergence time in million years ago (Mya).

**Supplementary table 1. List of primers**

| Name | Sequence |
|---|---|
| Oligonucleotides for CRISPR/Cas9 mutagenesis<br>* The underlined 4–bp sequence for ligation | |
| gRNA–1F | 5'– ATTGGTCTTCTTCGACATGACCAT –3' |
| gRNA–1R | 5'– AAACATGGTCATGTCGAAGAAGAC –3' |
| gRNA–2F | 5'– ATTGTCATGTCGAAGAAAACCTTA –3' |
| gRNA–2R | 5'– AAACTAAGGTTTTCTTCGACATGA –3' |
| gRNA–3F | 5'– ATTGCCATGACGATGCGACCGGCG –3' |
| gRNA–3R | 5'– AAACCGCCGGTCGCATCGTCATGG –3' |
| Primers for cloning of CDS of CyPA and RIN4<br>* The underlined sequences for pENTR/D–TOPO or BP reaction | |
| CyPA1–F | 5'– CACCATGTCTAACCCTAAGGTCTTCTTCG –3' |
| CyPA1–R | 5'– CTACGAAAGTTGACCGCAATCG –3' |
| RIN4–F | 5'– ACAAGTTTGTACAAAAAAGCAGGCT<br>ATGGCACAACGTTCTCATG –3' |
| RIN4–R | 5'– ACCACTTTGTACAAGAAAGCTGGGT<br>TCATTTCTTGCTAAACCCAAAG –3' |
| Site–directed mutagenesis by PrimeSTAR® Mutagenesis<br>* The underlined sequences for the mutation sites | |
| CyPA1–S58F–F | 5'– AAGGGCTTTTCCTTCCACCGTGTCATC –3' |
| CyPA1–S58F–R | 5'– GAAGGAAAAGCCCTTGTAGTGGAGAGG –3' |
| RIN4–P149V–F | 5'– GCTGTTGTGAAGTTTGGTGAGTGGGAC –3' |
| RIN4–P149V–R | 5'– AAACTTCACAACAGCAGCACCTTTCTC –3' |
| RIN4–ΔP149–F | 5'– GCTGTT__AAGTTTGGTGAGTGGGAC –3' |
| RIN4–ΔP149–R | 5'– AAACTT__AACAGCAGCACCTTTCTC –3' |
| Primers for Type III secretion system (T3SS) mutant<br>* The underlined 15–bp sequenced denote duplication for In–fusion | |
| T3SS–1F | 5'– CGGTACCCGGGGATCTGGAGCGAGTACGGCAATGT –3' |
| T3SS–1R | 5'– TTTCTGTCCTGGCTGGATTGAGCCACGCTCACTCAT –3' |
| T3SS–2F | 5'– AGTACCGCCACCTAATAAGAGGCAGCGCGATCGAA –3' |
| T3SS–2R | 5'– CGACTCTAGAGGATCGCACTAGCCCTCTTGTGTTA –3' |

| | | | |
|---|---|---|---|
| GmR–F | 5'– | CAGCCAGGACAGAAATGCCT | –3' |
| GmR–R | 5'– | TTAGGTGGCGGTACTTGGGT | –3' |
| Primers for qRT–PCR | | | |
| UBQ–F | 5'– | ATGCAGATCTTCGTCAAGACCTTG | –3' |
| UBQ–R | 5'– | ACCTCCCCTCAGACGAAG | –3' |
| RPS2–F | 5'– | GGGAGTTTCAAATGGTGGAAGA | –3' |
| RPS2–R | 5'– | CGGTTGTGCTGAAATTGGC | –3' |
| Chitinase–F | 5'– | TCAATTTGCTCGGTCAATGGG | –3' |
| Chitinase–R | 5'– | TCCCCATGCATTTGAAGCTT | –3' |
| Defensin–like–F | 5'– | CATGGTCAGGGCCTTGTTTT | –3' |
| Defensin–like–R | 5'– | CAAGCAAAACCAAAGCCCTG | –3' |
| ERN1-F | 5'– | TGGACATGCCTAAGACTGATGGC | –3' |
| ERN1-R | 5'– | TGAGCACAAGGGTGGAAGATCCCAC | –3' |
| NFR1–F | 5'– | CCCTTGTACCACAGAACC | –3' |
| NFR1–R | 5'– | GCTTTCTCTTCTTCCTTCTTCTG | –3' |
| RPG–F | 5'– | AAGGAGAACTTATTAGCAAGAGAA | –3' |
| RPG–R | 5'– | GTTGAATCTTGTCTCTCATTCATT | –3' |
